# vDeepInsight: an injective three-dimensional voxel carrier for tabular-feature neighborhood learning

**DOI:** 10.64898/2026.06.22.733711

**Authors:** Shangru Jia, Artem Lysenko, Keith A. Boroevich, Alok Sharma, Tatsuhiko Tsunoda

## Abstract

DeepInsight-style methods make tabular feature relationships accessible to convolutional networks by placing each feature at a fixed position on an image carrier. An open design question is how the carrier geometry should be constructed when feature neighborhoods themselves carry part of the signal. We introduce vDeepInsight, an injective three-dimensional (3D) voxel carrier that preserves feature neighborhoods more faithfully than matched two-dimensional (2D) carriers while keeping a one-to-one mapping from each feature to a single voxel. Genes are embedded with t-SNE or UMAP, assigned one-to-one to a sparse voxel grid by linear-sum assignment, and processed by a submanifold sparse 3D convolutional network. We evaluate the carrier on gene expression through four linked analyses. First, representation-quality metrics show that 3D layouts reduce gene-neighborhood distortion relative to matched 2D layouts before any model is trained. Second, controlled synthetic tasks show that a sparse 3D convolution can exploit this preserved locality, but only when the supervised signal is constructed to depend on co-located genes and the receptive field spans adjacent voxels. Third, on real omics tasks the 3D carrier matches or exceeds tuned tabular baselines and consistently exceeds matched 2D carriers; the margin is small on marker-type classification, where individual genes already carry much of the label (tissue, lineage and cancer-type classification), and larger on program-type tasks, where the target depends on coordinated, pathway-level multi-gene activity (drug-response regression, TCGA immunogenomic-context regression and mechanism-of-action classification). Fourth, because the assignment is injective, voxel attribution maps directly back to genes, enabling gene-level attribution and pathway-level functional interpretation without voxel-to-gene deconvolution. Overall, the added carrier dimension improves the fidelity of feature-neighborhood representation and translates this improvement into prediction gains that are largest when the signal is distributed across local gene programs rather than dominated by individual marker genes.

## 1 Introduction

Many prediction problems are presented to machine-learning models as tabular feature vectors whose entries are not exchangeable: the features have relationships that matter whenever the target depends on their joint behavior. DeepInsight introduced a general way to expose such structure to convolutional neural networks, by assigning each tabular feature to a fixed position on an image carrier and rendering every sample as an image; the construction is not specific to any one domain and has been applied across a range of tabular problems [1, 2, 3]. Gene expression is a clear instance of this setting, and the one used throughout this paper: genes participate in co-expression modules, pathways and regulatory programs, so that coordinated multi-gene activity, and not only individual marker genes, can carry the supervised signal. The same image-carrier principle has been used for single-cell annotation and multi-omics drug-response prediction [4, 5].

The methodological difficulty is that a carrier of this kind is not a natural image. Locality here means neighborhood in an unsupervised gene layout, not physical proximity. Translation equivariance and stationarity, which are well matched to photographs, have a weaker interpretation for a t-SNE or UMAP layout of genes. For this reason, the central design criterion is carrier fidelity: informative feature neighborhoods should remain close after layout fitting and discretization, and the downstream model should be able to combine those neighborhoods through its receptive field. This criterion matters because tuned gradient-boosted trees, linear models and tabular neural networks are strong baselines for medium-sized omics tables [6, 7, 8].

The established DeepInsight carrier is two-dimensional, and the question this study isolates is whether the same fixed-carrier principle preserves feature neighborhoods more faithfully when the carrier is expanded into three dimensions. The hypothesis is that an injective 3D voxel carrier improves neighborhood fidelity relative to matched 2D carriers, while keeping the one-to-one feature-to-voxel traceability that makes DeepInsight-style layouts interpretable. The evaluation has four parts, each aligned with one link in this argument. Representation-quality metrics test the premise directly: whether the 3D layout reduces neighborhood distortion before any model is trained. Synthetic neighborhood controls test the mechanism: whether a sparse 3D convolution can exploit a known local interaction when the signal is placed there by construction. Real benchmarks test the consequence: whether the same carrier stays competitive with the strong tabular baselines noted above on marker-type tasks, in which individual genes or compact expression modules already carry much of the label, and shows larger margins on program-type tasks, in which the target instead depends on coordinated, pathway-level multi-gene activity. Attribution analysis tests interpretability: whether the one-to-one voxel assignment supports gene-resolved biological readout.

vDeepInsight implements this design by embedding genes in ℝ^3^, assigning them to unique voxel centers by linear-sum assignment (LSA), and processing the occupied sites with a submanifold sparse 3D CNN (Figure 1). Matched 2D controls use either an independent DeepInsight-style 2D layout or a projected 2D layout derived from the same 3D coordinates. This makes the comparison primarily about carrier dimensionality and assignment geometry, rather than about unrelated preprocessing or model families.

**Figure 1:**
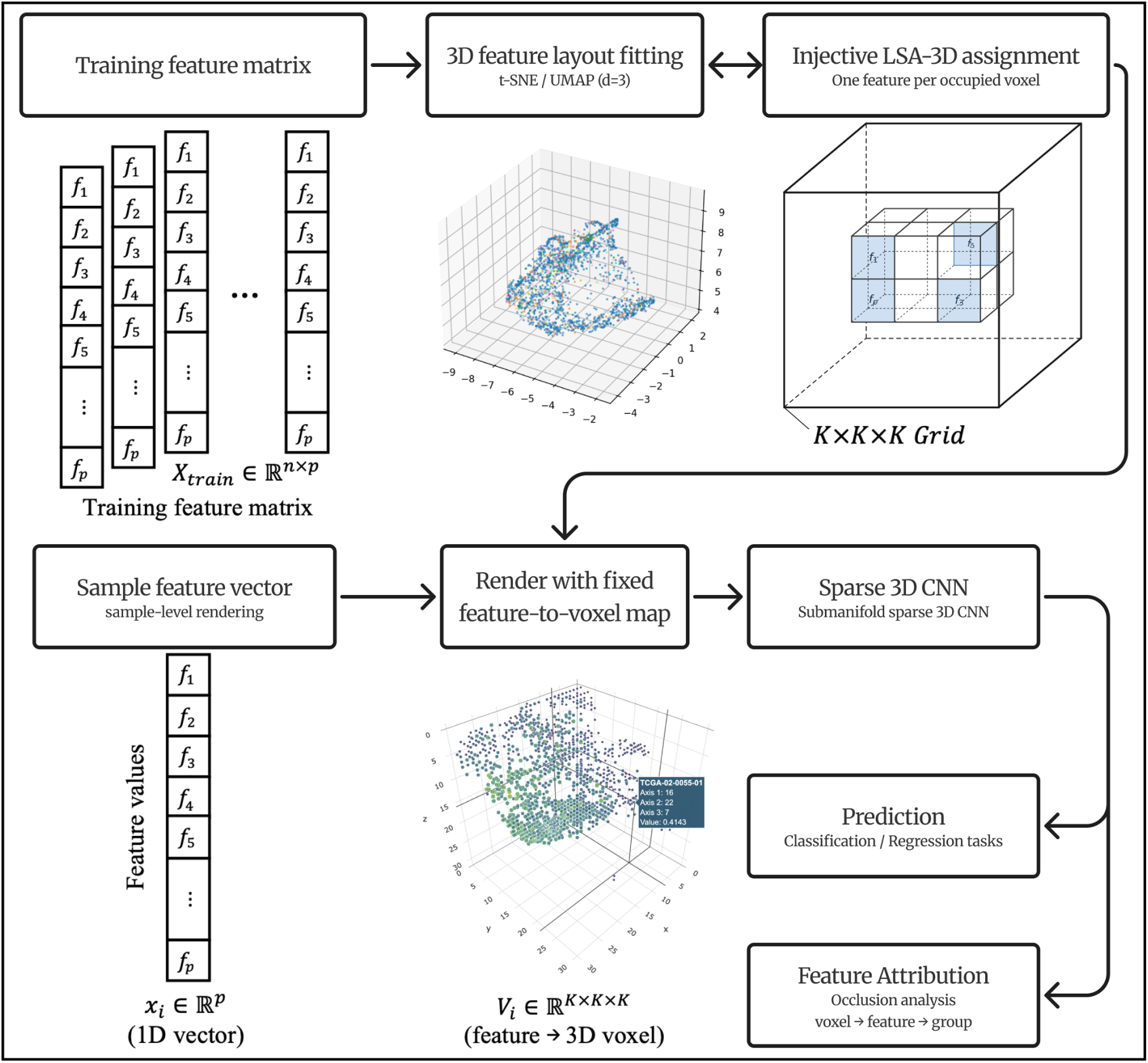
Overview of the vDeepInsight workflow. The figure summarizes the two-stage carrier construction and sample-rendering procedure. The carrier accepts any tabular feature set; gene expression is the running example used throughout this paper, so the schematic uses features *f_1_, . . ., f_p_* to denote genes or other tabular variables. *Top row:* a fixed feature-to-voxel map is fitted once from the training feature matrix *X*_train_ ∈ ℝ*^n^*^×^*^p^*. The *p* features are embedded in three dimensions (t-SNE or UMAP, *d*=3), and injective linear-sum assignment (LSA) assigns each feature to a unique occupied voxel, giving a one-feature-per-voxel layout. *Bottom row:* each sample feature vector *x_i_* ∈ ℝ*^p^* is rendered through this fixed feature-to-voxel map into a sparse voxel volume *V_i_* ∈ ℝ*^K^*^×^*^K^*^×^*^K^*. A submanifold sparse 3D convolutional network then processes the occupied voxels for classification or regression. Because the assignment is injective, voxel-level occlusion attribution maps directly back to features and then, for gene-expression data, to genes and pathways. Matched 2D carriers are used throughout the paper as controls under the same split, gene panel and preprocessing protocol.

Across representation metrics, synthetic controls and real omics benchmarks, vDeepInsight provides a consistent carrier-level advantage over matched 2D layouts. The downstream margin is smallest on near-saturated marker-type classification and largest on program-type prediction tasks, indicating that the added carrier dimension is most beneficial when local multi-gene structure contributes to the target.

## 2 Materials and methods

### 2.1 Study design

The study was organized as a matched carrier-dimension evaluation. Within each experimental block, the split, selected genes, preprocessing, labels and primary metric were fixed before changing the carrier or baseline model. This design makes the comparison primarily about representation and model class rather than about different data-processing paths. The evaluation contains four parts: representation-quality analysis, synthetic neighborhood controls, supervised benchmarks and attribution through the injective voxel map. Implementation details, code organization and baseline parameter choices are given in Supplementary Notes 1–2.

### 2.2 Datasets and preprocessing

The supervised datasets were selected to represent two signal regimes rather than to maximize the number of benchmarks. TCGA cancer type, GTEx tissue and CCLE/DepMap lineage classification are marker-type tasks in which individual genes or compact expression modules are expected to be informative. These datasets test whether the 3D carrier remains competitive when tabular baselines have limited headroom. GDSC drug response, TCGA immunogenomic-context regression and LINCS mechanism-of-action classification are program-type tasks in which the target is more likely to depend on coordinated pathway activity; these tasks test the setting in which a neighborhood carrier is expected to be more useful.

Bulk classification used TCGA pan-cancer RNA-seq data with 33 cancer-type classes [9, 10], GTEx tissue expression with 51 tissue classes [11], and CCLE/DepMap expression with 26 cell-line lineage classes [12]. Drug-response regression matched DepMap expression profiles to GDSC ln IC50 measurements for 676 drugs [13, 14]. MoA classification used LINCS L1000 landmark-gene signatures across 20 mechanism classes under a signature-level split [15]. TCGA immunogenomic-context regression used bulk TCGA expression and 17 continuous immunogenomic endpoints from the TCGA PanCancer immune-landscape resource of Thorsson et al. [16], covering immune-cell infiltration, neoantigen and mutation load, genome-integrity scores and repertoire-diversity measures. This task was evaluated on raw targets and on a within-cancer residualized panel.

Within each matched block, expression values were log-transformed where applicable and standardized using training-set statistics. Gene selection was also restricted to the training split. The bulk classification benchmarks used 2,000, 6,000 and 10,000-gene panels; GDSC used 6,000 genes; LINCS used the 978 L1000 landmark genes; and TCGA immunogenomic-context regression used 10,000 genes. The fitted layout, selected genes and scalers were then reused unchanged for validation and held-out test samples.

### 2.3 Three-dimensional voxel carrier construction

For a training expression matrix *X* ∈ ℝ^*n×p*^, genes were treated as samples by applying t-SNE or UMAP to *X*^⊤^ [17, 18]. This yields continuous coordinates

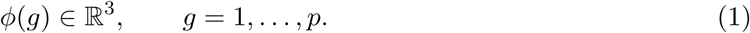

Coordinates were min–max scaled into grid space and assigned to unique voxel centers by solving an injective LSA problem,

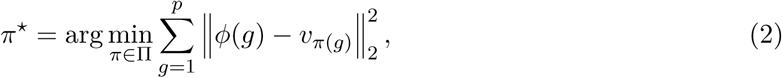

where {*v_j_*} are candidate voxel centers and Π is the set of injective gene-to-voxel maps. The resulting map is collision-free: each occupied voxel contains at most one gene, and every voxel-level attribution can be mapped back uniquely to the corresponding gene. For each sample *i*, the sparse volume *V* ^(i)^ is constructed by writing the standardized expression value 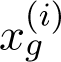 to voxel *π\**(*g*) and leaving all unassigned voxels empty.

The main bulk classification and immune experiments used a 32^3^ grid; GDSC drug response used a 24^3^ grid for the 6,000-gene panel; LINCS used a 12^3^ grid for the 978 landmark genes. This task-specific side length keeps occupancy in a practical range while preserving the same injective assignment principle. The representative 6,000-gene 32^3^ layout has occupancy approximately 18%.

### 2.4 Matched two-dimensional carriers and representation-quality metrics

Two 2D carriers were used to disentangle carrier dimensionality from other layout choices. The native DeepInsight-2D carrier fits an independent 2D t-SNE/UMAP layout and applies an analogous one-to-one 2D assignment. The Projected-2D carrier starts from the same 3D coordinates as vDeepInsight, applies a linear PCA(2) projection, and then solves the one-to-one 2D assignment. The latter isolates the effect of reducing the carrier dimension while holding the upstream coordinate construction as fixed as possible.

Representation quality was assessed on an identical 6,000-gene TCGA matrix using trustworthiness, continuity, kNN Jaccard overlap, Sammon-weighted stress and Kruskal-1 stress [19, 20, 21]. These metrics quantify different failure modes: trustworthiness penalizes false neighbors introduced by the embedding, continuity penalizes true neighbors lost by the embedding, kNN Jaccard reports direct neighborhood overlap, and stress measures global distance distortion.

The geometric expectation is intentionally weak but useful. For any stress-like objective, the space of 2D layouts is embedded in the space of 3D layouts by setting the third coordinate to zero; therefore

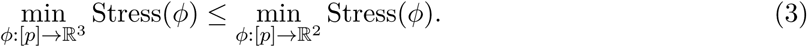

Equation 3 does not imply better prediction. It only motivates the representation-quality analysis: if a 3D carrier helps, it should first show a measurable reduction in neighborhood distortion.

### 2.5 Sparse 3D convolutional network architecture

Each rendered sample is a sparse third-order tensor in which only the occupied voxels—one per gene—carry a value and all remaining grid sites are empty. A dense *K*^3^ convolution over such a volume is wasteful, so the carrier is processed with a submanifold sparse convolutional network [22, 23], implemented with the spconv spatially sparse convolution library [24]. Two complementary operations are combined. Submanifold sparse convolutions compute outputs only at sites that are already active and keep the active-site pattern fixed, so they add depth and local feature mixing without dilating the sparse footprint or inflating computation; regular strided sparse convolutions are used at the transitions between resolution levels to coarsen the active set and enlarge the effective receptive field. Each convolution is followed by batch normalization and a ReLU nonlinearity.

The mainline backbone arranges these operations into a small number of resolution stages: a sparse stem first lifts the single-channel voxel value into a feature space, and alternating submanifold and strided sparse stages then progressively coarsen the grid while widening the receptive field. Because computation is restricted to occupied voxels, cost scales with the number of genes (the active sites) rather than with the full grid volume *K*^3^, which is what keeps the 3D carrier practical despite the added dimension.

The network produces a fixed-length descriptor by multi-scale mean–max pooling. At each retained resolution stage it applies global average pooling and global max pooling over the active voxels and concatenates these statistics across stages into a single vector (the readout referred to as ms_meanmax in the implementation). Average pooling summarizes the typical activation of each feature channel while max pooling captures its strongest local response, and aggregating both across scales retains signal that is either spatially diffuse or locally concentrated in the gene neighborhood. The pooled descriptor is passed to a lightweight multilayer-perceptron head for classification or regression. A deeper residual sparse-3D variant was also implemented and evaluated as an ablation; it did not give a consistent improvement over this plain multi-scale mean–max backbone, including on the LINCS and immunogenomic-context tasks, so the plain backbone is used for all headline comparisons unless explicitly stated. Exact channel widths, stage counts, optimization settings and task-specific loss functions are listed in Supplementary Note 2.

### 2.6 Synthetic neighborhood controls

The synthetic experiments were designed as a positive capability test in which local gene co-location is required by construction. They provide a controlled setting for testing whether the sparse 3D architecture can read neighborhood-dependent signals through the voxel carrier. The central design choice is that the gene-expression values themselves carry no usable marginal signal. Starting from a real layout artifact, we retained only its grid geometry and train/validation/test split sizes and overwrote all voxel features with independent standard-Gaussian gene values 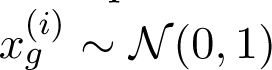 ∼ *N* (0, 1). The ground-truth label was then derived from these synthetic values. Because the same synthetic gene-by-sample matrix and the same labels are reused for every layout, and only the gene-to-voxel placement changes, any performance difference between layouts is attributable to placement geometry rather than to the data. This is a deliberate departure from planting labels on real expression values, which would let pointwise or histogram shortcuts solve the task before any neighborhood was used.

To make the co-located gene set reproducible and to align it with the 3^3^ receptive field used below, the patch *P* was defined as a single 3 × 3 × 3 voxel window of the real layout. We centered the window on the occupied voxel whose surrounding 3 × 3 × 3 neighborhood contained the largest number of occupied voxels (the densest local neighborhood; ties broken by raster voxel index), and took *P* to be the set of genes occupying that window (at most 27 genes, and fewer in practice because the carrier is sparse). The gene at the central voxel was designated the anchor *a*. By construction every gene in *P* lies within a single 3^3^ kernel footprint, so one convolution centered on the window can read all of *P* at once, whereas position-blind or 1^3^ receptive fields cannot. Writing 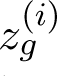 for the training-standardized value of gene *g* in sample *i*, we defined three signal scores of increasing reliance on co-location:

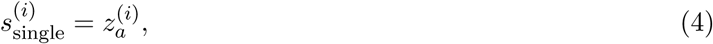

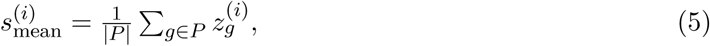

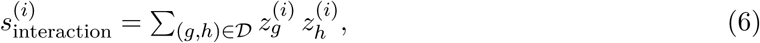

where *D* is a set of disjoint gene pairs drawn from *P*. Each score was converted to a balanced class label by thresholding at its training-set median. The single-gene score (Equation 4) depends on one marker and should be solvable by any position-aware readout. The patch-mean score (Equation 5) is additive and can in principle be solved by reading patch genes separately and summing them, so it does not strictly require co-location. The patch-interaction score (Equation 6) has uninformative single-gene marginals—each *z_g_* is iid Gaussian, so no individual gene predicts the label—and places all signal in pairwise products; it can therefore only be read by a model that brings interacting genes into a shared receptive field.

Two factors were then crossed. The *layout* was either the real structure-preserving t-SNE/UMAP placement, a random-coordinate placement, or a fully randomized layout; the latter two scatter the patch genes across the grid. The *receptive field* was one of three settings: position-blind global pooling (a 1^3^ filter with no downsampling followed by global pooling), a 1^3^ strided stack that is position-aware but cannot mix neighboring voxels, or a local 3^3^ strided stack that can combine adjacent voxels. Crossing three signals, three layouts and three receptive fields gives the 27 conditions reported below. The decisive condition is the patch-interaction signal under the real layout with a 3^3^ receptive field: only there are the interacting genes both co-located by the carrier and jointly visible to a single kernel.

### 2.7 Training, baselines and statistical testing

All supervised comparisons were run inside the matched blocks defined above. The 3D and 2D neural carriers used the same split and gene panel within a block, and tabular baselines operated on the corresponding standardized expression matrix. The classification baselines were random forests [25], histogram gradient boosting [26], XGBoost [27] and the FT-Transformer [28]; the regression tasks additionally used a ridge linear baseline (GDSC drug response), and the LINCS task used a multilayer perceptron on the raw signature. The exact baseline set for each task is summarized in Supplementary Note 2.

Macro-F1 was the primary metric for classification because class frequencies are imbalanced. Accuracy is reported as a secondary metric. GDSC regression used per-drug Pearson and Spearman correlation on held-out cell lines. TCGA immunogenomic-context regression used per-target Pearson correlation on held-out samples; win/tie/loss against random forests used a ±0.01 practical-equivalence margin. Paired Wilcoxon signed-rank tests and paired bootstrap confidence intervals were used for matched comparisons, with Benjamini–Hochberg false-discovery-rate control across the main classification comparisons [29].

### 2.8 Attribution and pathway enrichment

Attribution was computed by occluding occupied voxels in trained TCGA models, averaging importance across model seeds and mapping each occupied voxel back to its unique gene. The resulting seed-averaged gene rankings were summarized by pre-ranked enrichment against KEGG pathways, providing pathway-level functional readouts of the genes used by the model. Cross-class distinctness was quantified by the pairwise Jaccard overlap of per-class top-attributed gene sets. This analysis depends on the injective voxel assignment: no post-hoc deconvolution is required to decide which gene an attributed voxel represents.

## 3 Results

The results follow the four-part argument set out in Section 2.1. Sections 3.1–3.3 examine the carrier without using any real task labels: the injective assignment and the neighborhood it preserves are measured directly on the gene layout (Sections 3.1–3.2), and the synthetic controls use labels constructed by design rather than observed outcomes (Section 3.3). Sections 3.4–3.7 then report performance on the real supervised tasks, kept separate by signal regime: marker-type classification (Section 3.4) and program-type regression and classification (Sections 3.5–3.7). Sections 3.8 and 3.9 report training cost and gene-level attribution. Separating the representation analysis from the supervised analysis is deliberate: it lets us state precisely what the 3D carrier changes about the representation before asking, task by task, whether that change improves prediction.

### 3.1 Injective voxel assignment preserves gene traceability

The vDeepInsight carrier constructs a sparse volume in which each occupied voxel corresponds to exactly one gene. This injective design differs from non-injective rasterization schemes: there is no collision averaging, and therefore no ambiguity in attribution. The representative 32^3^ layouts used in the bulk experiments operate at approximately 18% occupancy for 6,000 genes, leaving enough empty space for the LSA step to preserve the continuous gene neighborhood while keeping the grid sparse enough for efficient submanifold convolutions. A separate LSA diagnostic confirmed that discretization itself is not the limiting step: at a first-layer CNN scale of *k* = 26, continuous-to-voxel assignment retained trustworthiness and continuity near 0.999, and 90.6% of continuous neighbors fell within a 5 × 5 × 5 voxel window.

### 3.2 Three-dimensional layouts reduce gene-neighborhood distortion

The representation-quality analysis was consistent with the geometric expectation (Figure 2). Global stress was markedly lower for 3D carriers than for 2D carriers: Sammon stress was 45.2 for 3D t-SNE and 53.6 for 3D UMAP, compared with 68.1–70.9 for 2D layouts; Kruskal-1 stress was 0.221 and 0.234 for 3D t-SNE and UMAP, versus 0.265–0.269 for 2D layouts. The 3D t-SNE carrier also had the highest local preservation at small neighborhood sizes, including trustworthiness *T* (5) = 0.979. The dominant axis of variation across the five carriers is the embedding algorithm (t-SNE versus UMAP) rather than the carrier dimension, so the appropriate test is the matched within-algorithm comparison. Holding the embedding algorithm fixed, each 3D layout improved on its 2D counterpart on every metric—3D t-SNE over native 2D t-SNE, and 3D UMAP over 2D UMAP, for trustworthiness, continuity, kNN Jaccard, Sammon stress and Kruskal-1 stress; the global-stress reduction is the cleaner cross-carrier signal, holding for both 3D variants against all 2D layouts. These values instantiate Equation 3: the third dimension reduces flattening distortion before any prediction model is trained.

**Figure 2:**
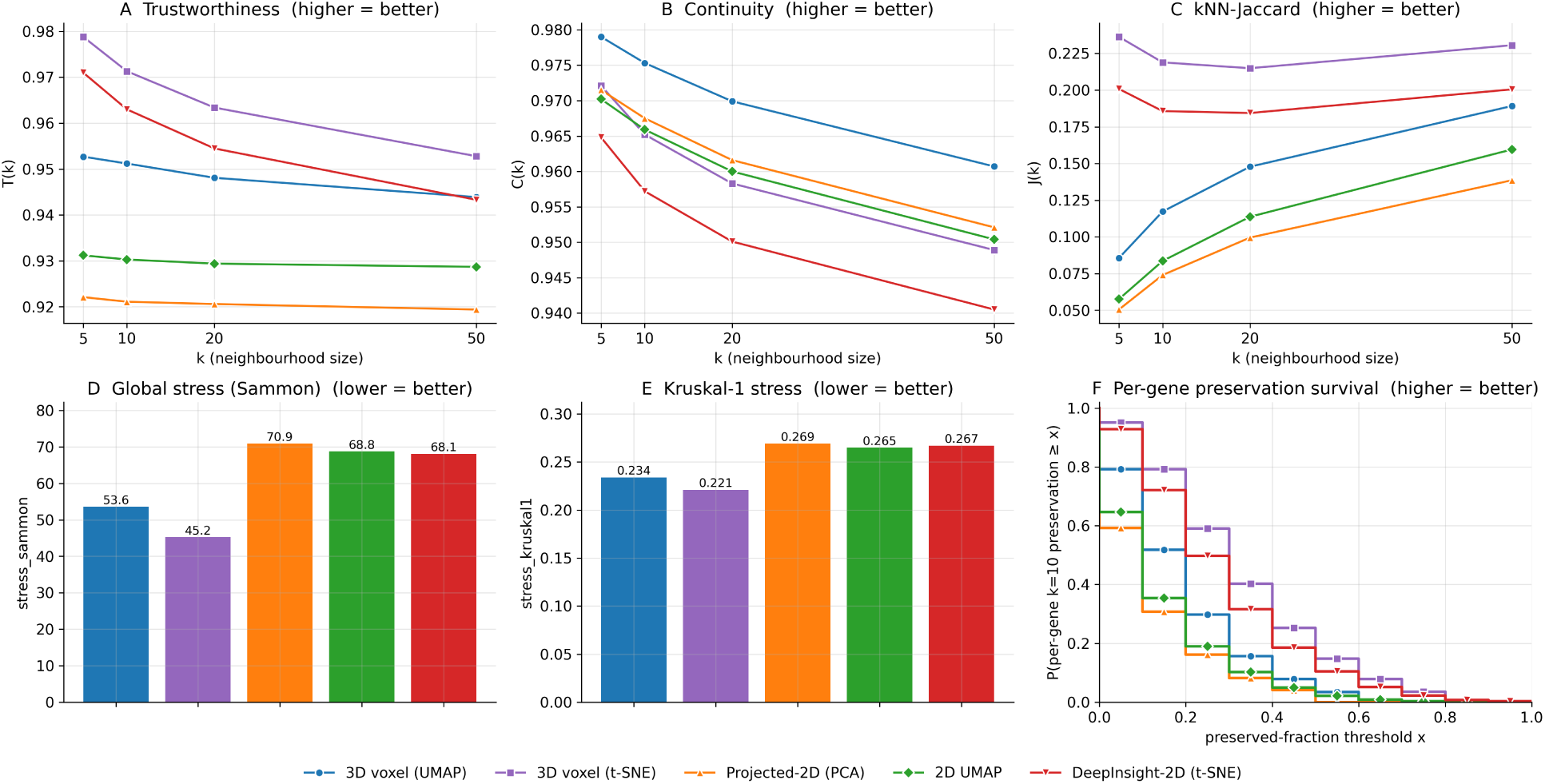
The 3D layout reduces gene-neighborhood distortion. Five carriers are compared on an identical 6,000-gene TCGA matrix. **(A–C)** Trustworthiness, continuity and kNN Jaccard as functions of neighborhood size *k*. **(D–E)** Sammon-weighted and Kruskal-1 stress; lower values indicate less global distortion. **(F)** Per-gene *k* = 10 preservation survival curve. Both 3D variants reduce global stress relative to all 2D layouts, and 3D t-SNE also gives the highest small-*k* local preservation.

Importantly, this is a representation result rather than a performance claim. Lower distortion means that a local 3D kernel has better access to the gene neighbors defined by the unsupervised layout. Whether that access matters depends on the signal in the downstream task.

### 3.3 Synthetic controls verify local-neighborhood exploitability

Following the design in Section 2.6, the three control patterns appear exactly as constructed (Figure 3). The position-blind global-pooling readout stayed at chance for every signal and layout (macro-F1 0.34–0.42), confirming that the labels cannot be solved without spatial information. The single-gene marker was recovered by any position-aware model irrespective of layout (macro-F1 0.95–0.99), and the additive patch-mean signal was largely layout-robust (macro-F1 0.94–0.97 on the real layout versus 0.83–0.90 on the randomized layouts), as expected for signals that do not require gene co-location.

**Figure 3:**
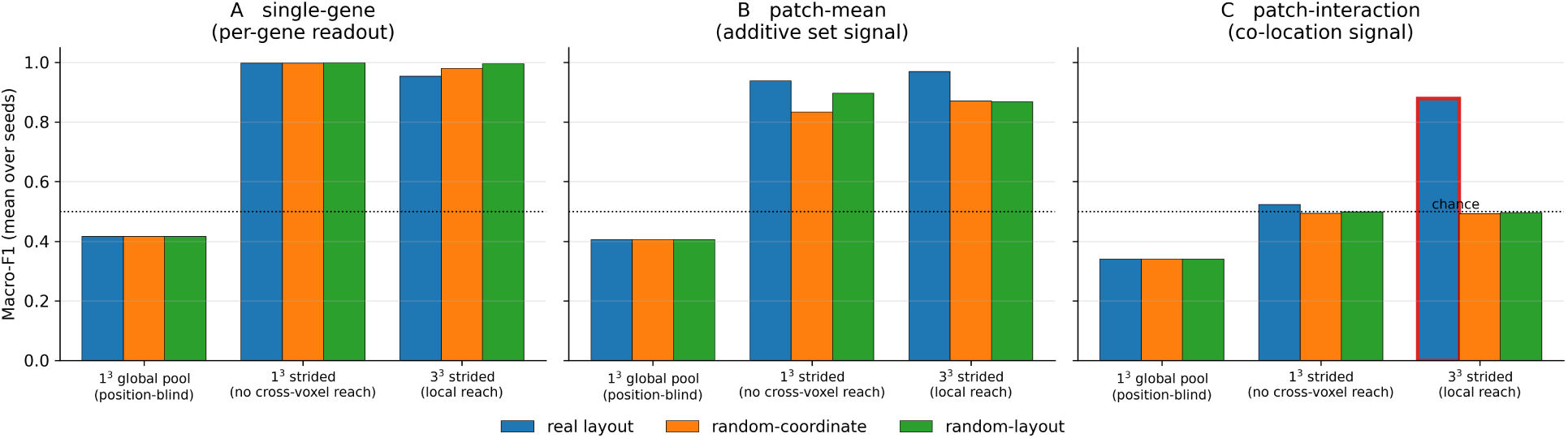
A sparse 3D CNN exploits neighborhood only when the signal lives there. Macro-F1 for three synthetic signal types crossed with three layouts and three receptive-field settings. A single-gene signal is layout-invariant once the readout is position-aware; an additive patch signal is largely robust to layout changes; the co-location interaction is recovered only with the real layout and a local 3^3^ receptive field. Randomization is used here only as a controlled capability probe.

Decisively, the co-location interaction was recovered only when both ingredients were present. With the real layout and a 3^3^ receptive field the model reached macro-F1 0.879, whereas removing local reach (real layout, 1^3^ stack) collapsed performance to 0.523 and removing co-location (random-coordinate or random-layout placement with the 3^3^ stack) collapsed it to 0.492 and 0.496—all three at chance. The interaction signal therefore behaves exactly as constructed: a sparse 3D CNN can read a planted pairwise-product signal only if the carrier places the interacting genes together *and* the kernel spans adjacent voxels, and neither ingredient suffices alone.

This experiment establishes the capability link between representation fidelity and downstream prediction. When a supervised signal is constructed to depend on co-located genes, the sparse 3D CNN can exploit the preserved neighborhood. The result connects the representation advantage of Section 3.2 to the real-task analyses below by showing that the architecture provides the required local readout mechanism rather than only a higher-capacity classifier.

### 3.4 Marker-type classification shows stable performance across gene scales

On GTEx, CCLE and TCGA classification, the 3D carrier had the highest mean macro-F1 across matched blocks (Figure 4). It achieved mean macro-F1 0.959 on GTEx, 0.754 on CCLE and 0.943 on TCGA. Against tree, transformer and 2D-carrier baselines, paired comparisons were significant after Benjamini–Hochberg correction in the main classification comparison. Of the two 2D carriers, native DeepInsight-2D is the matched image-carrier comparator under this controlled protocol; Projected-2D (a linear PCA(2) of the same 3D coordinates) is included only as a dimensionality-reduction ablation that isolates the effect of removing the third axis, and is not intended as a competitive 2D construction.

**Figure 4:**
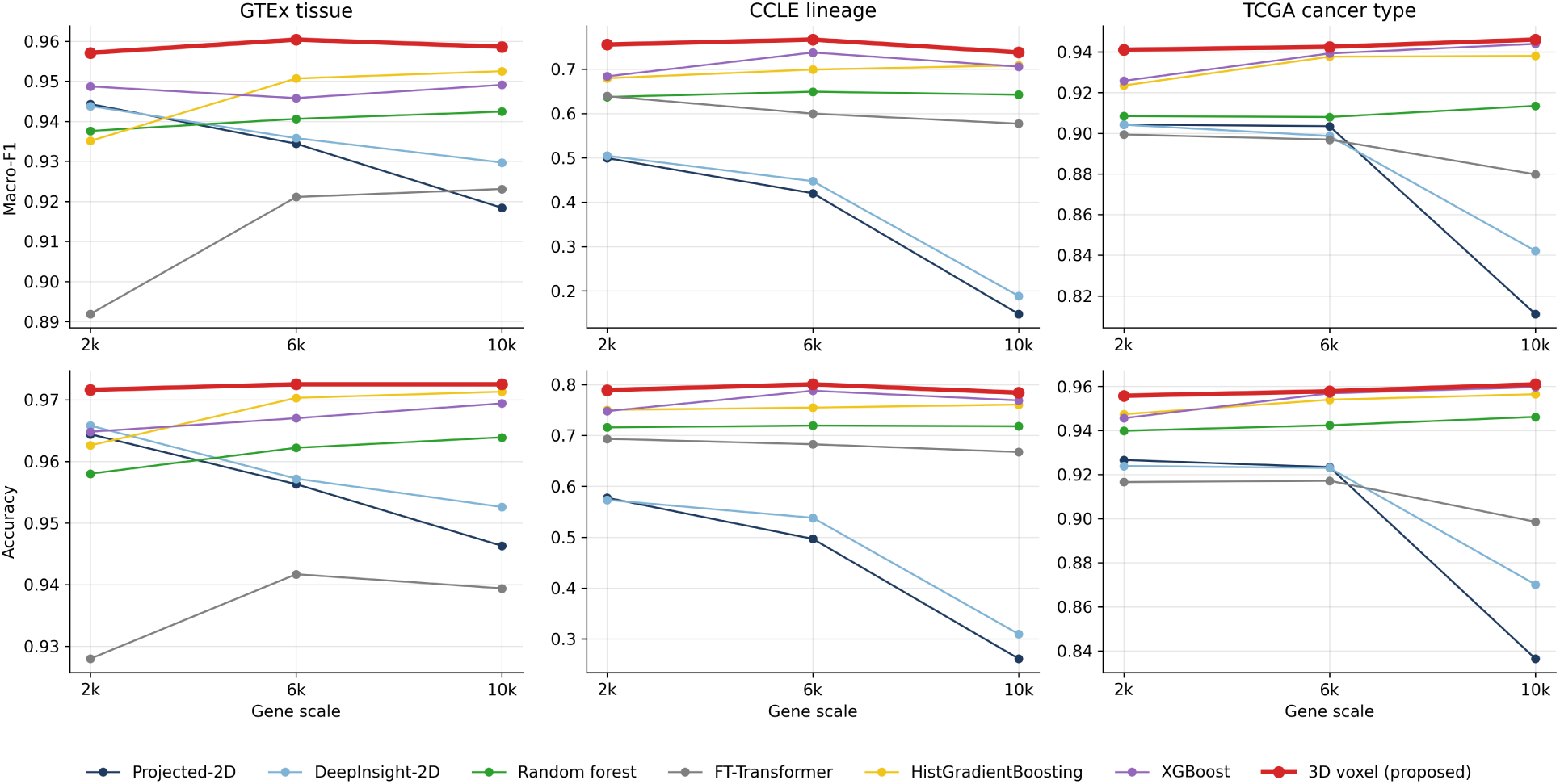
Classification across gene scales. Macro-F1 and accuracy for GTEx tissue, CCLE lineage and TCGA cancer-type classification as the gene panel grows from 2,000 to 6,000 and 10,000 genes. The 3D voxel carrier has the highest mean score on each dataset under this protocol and remains stable as gene count increases; the native 2D and Projected-2D carriers degrade most strongly on CCLE.

The size of the margin differed across datasets, which is consistent with the expected task regimes. GTEx tissue and TCGA cancer type are near-saturated, marker-dominated classification tasks. There the improvement from the 3D carrier over the strongest tabular baseline (XGBoost) was modest: approximately +0.011 on GTEx and +0.007 on TCGA. This is the expected low-end regime for a neighborhood carrier: marker genes already encode much of the label. CCLE showed a larger gap: +0.044 over the strongest tabular baseline, and approximately +0.37 over the 2D carriers. Because the LSA assignment is injective in both 2D and 3D—each gene occupies its own cell, with no pixel collisions—this 2D degradation is not caused by multiple genes crowding into shared pixels. The matched 2D carriers were in fact below the tabular baselines at every gene scale and fell further as the panel grew, which is consistent with reduced neighborhood fidelity of the flattened 2D layout together with a fixed-capacity matched 2D CNN, rather than with grid capacity. We therefore read the credible CCLE margin as the +0.044 improvement over tuned tabular baselines, and the much larger gap over the 2D carriers as a scale-stability contrast: 3D layouts remain more scale-stable than a matched 2D image carrier as the gene count grows.

### 3.5 Drug-response regression gives the largest observed real-task margin

GDSC drug-response regression gave the largest observed real-task margin for the carrier (Figure 5A). The 3D voxel model achieved per-drug Pearson *r* 0.470 ± 0.015 and Spearman *ρ* 0.451 ± 0.003, compared with 0.446/0.421 for histogram gradient boosting, 0.435/0.417 for ridge regression and 0.389/0.367 for the Projected-2D carrier. For GDSC the only 2D comparator is the Projected-2D dimensionality ablation (native DeepInsight-2D was not run on this regression task), so the most informative GDSC contrasts are against the tuned tabular baselines, in particular histogram gradient boosting (a gradient-boosted tree ensemble), confirming that the carrier’s advantage does not depend on the weakness of the projected 2D carrier. The improvement is concentrated rather than uniform: the ten drugs with the largest gain over histogram gradient boosting (Figure 5B) reach ΔPearson of +0.16 to +0.31—for example fulvestrant (+0.31), erlotinib (+0.27) and saracatinib (+0.23)—each lifting a drug that the tree baseline predicts poorly to a substantially higher held-out correlation.

**Figure 5:**
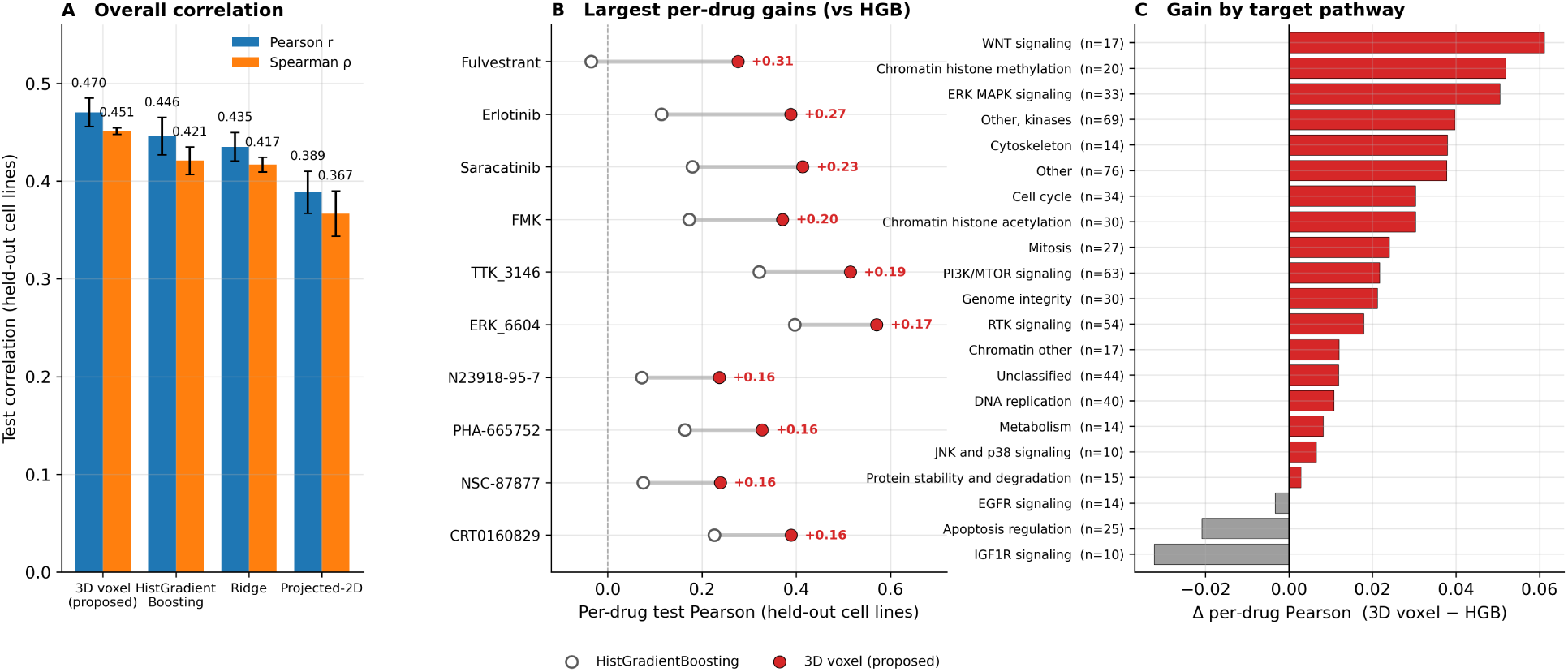
GDSC drug-response regression: where the 3D carrier helps most. DepMap expression was matched to GDSC ln IC50 measurements for 676 drugs over three cell-line-disjoint splits. **(A)** Overall per-drug Pearson and Spearman correlation by method (3D voxel, histogram gradient boosting, ridge and the Projected-2D dimensionality ablation; mean ± SD over splits). **(B)** The ten drugs with the largest per-drug gain of the 3D carrier over the strongest baseline (histogram gradient boosting); each row shows the held-out Pearson of histogram gradient boosting (open) and of the 3D carrier (filled), with the gain in Pearson annotated. Drugs are averaged over the three splits before differencing. **(C)** Mean per-drug ΔPearson (3D voxel minus histogram gradient boosting) within each drug-target pathway (pathways with at least ten drugs; bars colored by sign, drug count annotated). Gains concentrate on multi-gene-program pathways and are near zero or slightly negative on a few single-target pathways. All panels are computed directly from the recorded per-drug result tables and the GDSC drug-target annotation.

Resolving these gains by drug-target pathway shows a broadly positive pattern (Figure 5C). The per-drug advantage over histogram gradient boosting is positive for most pathways and is somewhat larger on multi-gene-program classes—including WNT signaling, chromatin histone methylation and ERK–MAPK signaling—than on single-target classes, with a mean ΔPearson of +0.032 across program-type pathways versus +0.014 elsewhere; a few single-target pathways (for example EGFR, IGF1R and p53) are near zero or slightly negative. We interpret this as a hypothesis-consistent descriptive observation, not as a standalone predictor of when the method will work. The more robust conclusion is the task-level result: a relational drug-response problem shows a wider and more broadly distributed 3D advantage than marker-type classification.

### 3.6 TCGA immunogenomic-context regression retains the carrier margin after within-cancer residualization

The TCGA immunogenomic-context regression provides a second program-type regression test (Figure 6). The 17 endpoints include estimates of immune-cell infiltration (for example leukocyte and lymphocyte fractions), neoantigen and somatic-mutation load, genome-integrity and aneuploidy scores, and T- and B-cell repertoire-diversity measures. Each target was predicted from bulk TCGA expression (10,000 genes) under the same matched protocol used for the other tasks, with per-target held-out Pearson correlation as the endpoint and random forests as the strongest tabular baseline in this experiment. The 3D carrier was numerically higher than random forests on 14 of 17 targets; under a ±0.01 practical-equivalence margin this corresponds to 11 wins, 6 ties and no losses, with a median per-target gain of +0.022 (Wilcoxon *p <* 0.001).

**Figure 6:**
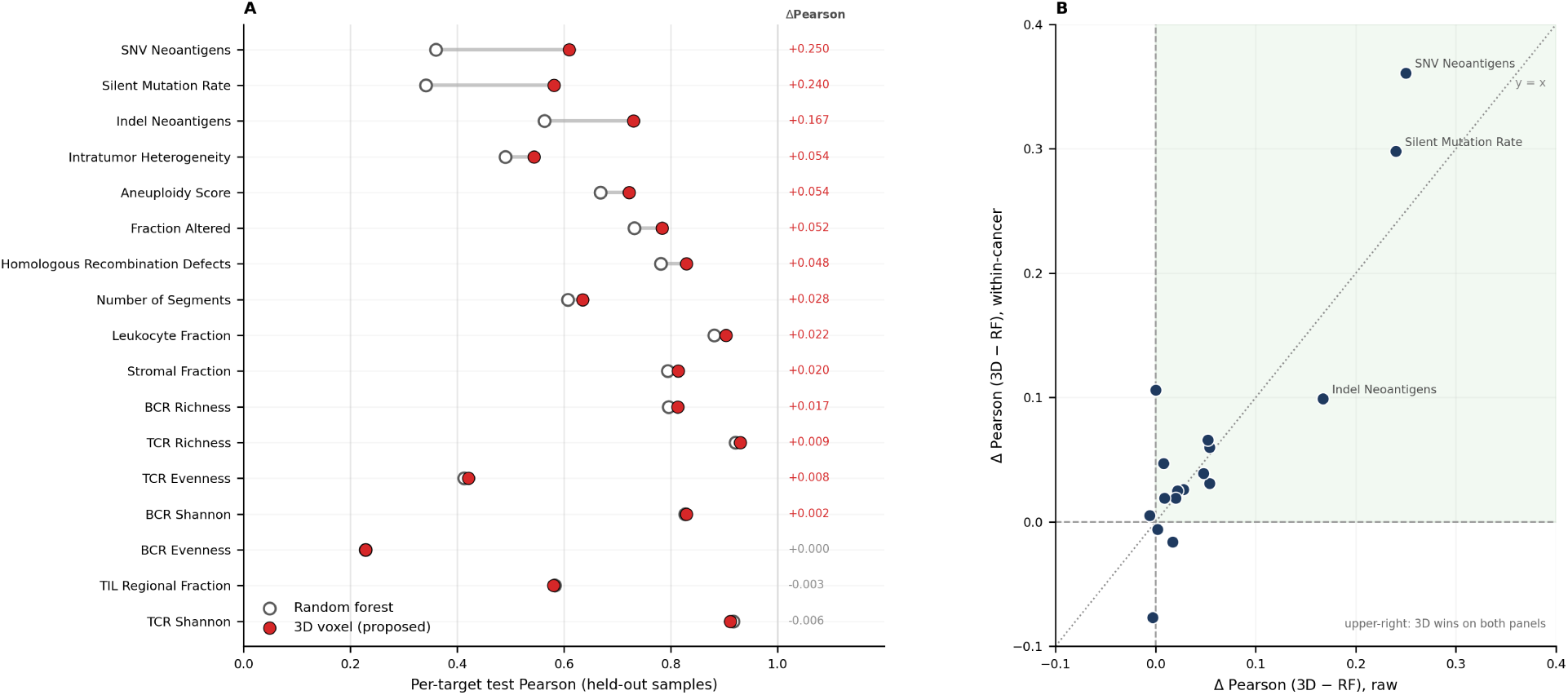
TCGA immunogenomic-context regression. **(A)** Per-target held-out Pearson correlation for the 3D voxel carrier and random forests, ordered by 3*D* − RF gain; the right column lists per-target ΔPearson. Using a ±0.01 practical-equivalence margin, 3D had 11 wins, 6 ties and 0 losses against random forests. **(B)** Per-target ΔPearson against random forests on raw targets versus within-cancer residualized targets. Most targets fall in the region where 3D wins on both panels, reducing the likelihood that the raw-panel result is explained only by cancer-type differences.

These raw immune endpoints are not independent of tumor lineage: several of them, such as leukocyte fraction, vary systematically across cancer types, so a model could in principle achieve part of its accuracy simply by reading cancer-type identity from expression rather than by capturing within-tumor immune signal. The within-cancer residualized panel was designed to remove exactly this confound. Each target was residualized against cancer type—subtracting the per-cancer-type mean so that only within-cancer-type variation remains—and all models were retrained and re-evaluated on these residuals, which isolates predictive signal that cannot be explained by cancer-type identity alone. After residualization the 3D advantage persisted and was slightly larger, with a median within-cancer gain of +0.031 compared with +0.022 on the raw panel (Figure 6B). This does not establish a causal mechanism, but it makes the cancer-type-shortcut explanation unlikely and is consistent with the program-type hypothesis: the carrier’s benefit on this task reflects distributed, within-cancer multi-gene signal rather than coarse differences between tumor lineages.

### 3.7 LINCS mechanism-of-action classification provides an additional program-type evaluation

LINCS mechanism-of-action classification provided an additional transcriptional-program task. Under a signature-level split across 20 MoA classes, the 3D carrier achieved macro-F1 0.577 ± 0.021 and accuracy 0.611 ± 0.010 (Figure S1). This exceeded an MLP on the raw signature (0.561/0.603), random forests (0.487/0.552), Projected-2D (0.437 macro-F1) and native DeepInsight-2D (0.428 macro-F1). The margin over the MLP is modest, but the result is consistent with the broader pattern: program-type tasks show a useful 3D advantage, while the 2D carriers are substantially weaker in this setting.

### 3.8 Sparse 3D convolution provides a practical accuracy–compute trade-off

Across the compact task summary (Table 2), the 3D carrier was competitive with the strongest recorded baseline in each setting. Comparisons with the matched 2D carriers are reported in the task-specific figures and text rather than pooled into a single column, because the evaluation units differ across tasks. Its computational profile was also practical in these matched runs (Figure 7). Sparse convolutions restrict computation to occupied voxels, so the 3D model is not simply a dense *K*^3^ CNN. At the 10,000-gene scale on TCGA, the 3D carrier trained in roughly 660 seconds, compared with approximately 1,100 seconds for XGBoost and 1,580 seconds for histogram gradient boosting. All reported training times were measured on a single server equipped with 4× NVIDIA A100 80GB GPUs. Random forests and FT-Transformer variants can be faster, but they occupy lower-accuracy regions in these matched comparisons. The result is not overall dominance; it is a practical accuracy–compute trade-off for tasks where the representation advantage matters.

**Figure 7:**
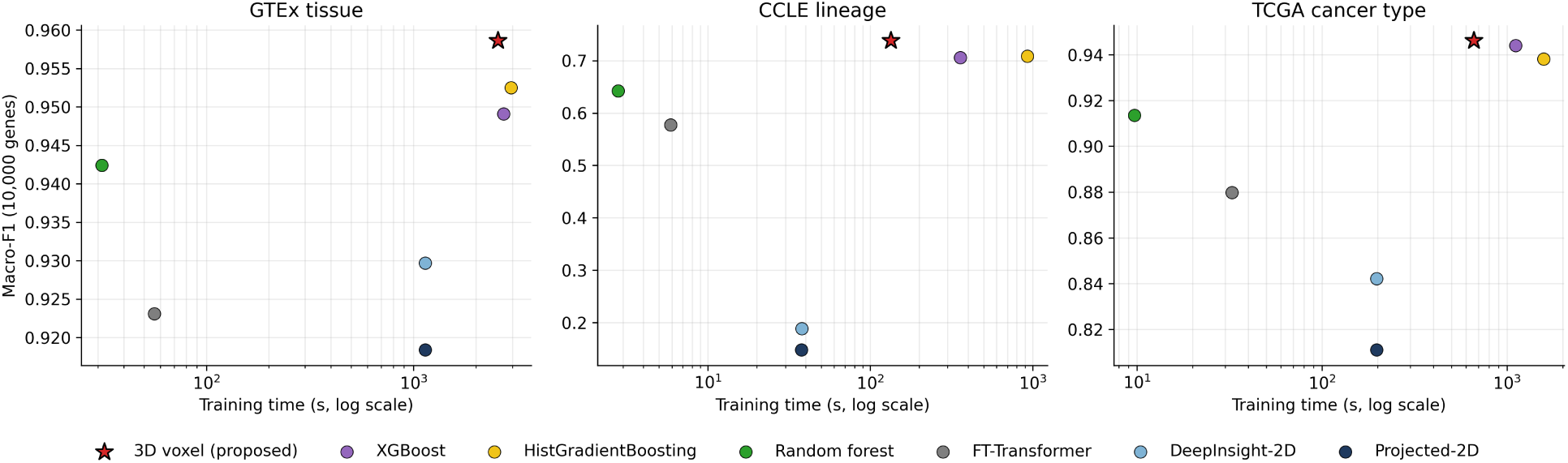
Accuracy versus training time. Macro-F1 against log-scaled training time for GTEx, CCLE and TCGA. The sparse 3D voxel carrier occupies a high-accuracy region in these matched comparisons; at the largest gene scale it trains faster than the gradient-boosted tree ensembles whereas some simpler baselines are faster but have lower accuracy under the same protocol. Training times were measured on a server with 4× NVIDIA A100 80GB GPUs.

### 3.9 One-to-one voxel attribution yields gene-resolved pathway interpretation

Because the voxel assignment is injective, attribution over occupied voxels maps directly back to genes without collision deconvolution (Figure 8). In the TCGA attribution analysis, voxel-occlusion scores were averaged across model seeds before gene ranking, providing a seed-aggregated view of model importance. This design preserves the main interpretability advantage of the carrier: each attributed voxel corresponds to a single gene, and each gene can be linked directly to downstream pathway-level functional summaries.

**Figure 8:**
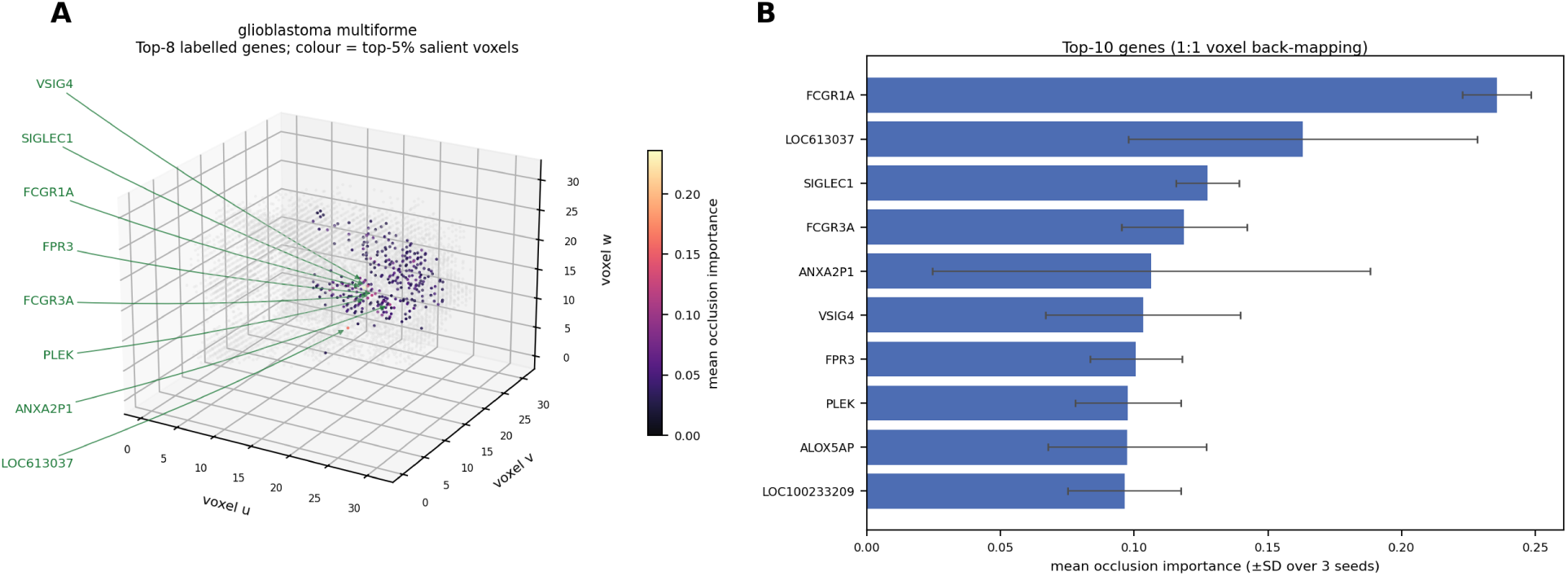
Gene-resolved attribution from the one-to-one voxel map. A single-class example is shown for glioblastoma multiforme in TCGA. Voxel-level occlusion attribution maps uniquely back to genes because every occupied voxel corresponds to exactly one gene. Seed-averaged attribution highlights an immune/myeloid program, including *FCGR1A*, *SIGLEC1*, *VSIG4* and *FCGR3A*. Pan-class attribution maps and pathway-level functional summaries are shown in Supplementary Figures S2 and S3.

In the glioblastoma example, the leading attributed genes included *FCGR1A*, *SIGLEC1*, *VSIG4* and *FCGR3A*, forming an immune/myeloid program consistent with known myeloid infiltration in this tumor type. The same attribution workflow was then applied across TCGA cancer classes. The per-class attribution overview (Supplementary Figure S2) shows that attribution is spatially concentrated in class-specific regions of the layout rather than diffusely spread across the full carrier. Related cancer groups, including GBM/LGG, LUAD/LUSC and KIRC/KIRP, show distinct attribution patterns, indicating that the model uses class-specific gene programs rather than a single generic pan-cancer signal.

Pathway enrichment was used as a functional aggregation of the seed-averaged gene attribution rankings (Supplementary Figure S3). These pathway-level summaries provide biological readouts of the genes used by the trained model and connect voxel-level occlusion importance to interpretable gene programs. Per-class top-gene sets were also largely distinct across cancer types (Supplementary Figures S4 and S5), further supporting class-specific use of the feature layout. This analysis should be viewed as model interpretation rather than causal perturbation evidence, but it demonstrates that the injective 3D carrier supports direct gene- and pathway-level interpretation in addition to prediction.

## 4 Discussion

### 4.1 Principal findings

This study evaluates an injective 3D voxel carrier as a carrier-design extension of the DeepInsight family of tabular-to-image methods, demonstrated here on gene expression. The results support three linked findings. First, the 3D layout reduces gene-neighborhood distortion relative to matched 2D layouts before any supervised model is trained. Second, a sparse 3D CNN can use that preserved neighborhood in a controlled setting when the label depends on co-located genes and the receptive field spans adjacent voxels. Third, in real tasks the 3D carrier matches or exceeds the strongest tuned tabular baseline on every evaluated task and consistently exceeds the matched 2D carriers; the size of the gain varies with task structure, from small on near-saturated marker-type classification to larger on drug-response and immunogenomic-context regression where distributed multi-gene signal is more plausible. The method is best interpreted as a geometry-aware extension of DeepInsight: it preserves feature neighborhoods more faithfully than matched 2D carriers and gives the largest gains when those neighborhoods encode distributed biological programs.

### 4.2 Methodological interpretation

The central methodological point is the separation of representation fidelity from downstream accuracy. Equation 3 gives only a feasibility argument: a 2D layout can be embedded in 3D space, so the optimal 3D stress cannot be worse under the same objective. The empirical quality analysis shows that this expected reduction is measurable for the gene panels used here. However, lower distortion has no automatic implication for supervised prediction. It becomes useful only if the preserved neighbors carry task-relevant joint signal and if the model architecture can combine them.

The synthetic controls were designed to test this second condition without relying on uncertain biological interpretation. The co-location interaction was recovered only when the structure-preserving (real) layout was combined with a receptive field that spanned adjacent voxels. The same experiment also explains why downstream margins depend on signal structure: single-gene and additive patch signals can be solved without depending on the specific placement of genes in the carrier, whereas interaction signals benefit from preserved local neighborhoods. This explains the real-task pattern. TCGA and GTEx are marker-dominated classification problems, in which individual genes already account for much of the label, so the improvement from the 3D carrier over tuned tabular baselines is small. GDSC drug response and the TCGA immunogenomic-context targets are more compatible with distributed, pathway-level signal, and show larger observed margins under the current protocols.

### 4.3 Relationship to 2D DeepInsight, tabular baselines and biological graphs

Building on DeepInsight, vDeepInsight extends the fixed spatial carrier from a 2D image to an injective sparse 3D voxel grid, improving feature-neighborhood preservation while retaining direct feature-level traceability. It is therefore complementary to the established 2D DeepInsight family rather than a competitor to it: the 2D route has practical advantages, including mature image backbones and transfer-learning infrastructure, while the 3D route offers a more faithful local layout and a collision-free mapping from occupied carrier sites back to genes. The present comparison holds the gene panel, split and preprocessing fixed so that the effect being tested is primarily carrier dimensionality and assignment geometry.

The strength of the tabular baselines also bounds how large the carrier’s advantage can appear. Gradient-boosted trees, random forests, linear models and tabular neural networks are appropriate competitors for expression matrices. The small margins on GTEx and TCGA are therefore not a weakness of the evaluation; they indicate that a neighborhood carrier has limited headroom when markers already solve much of the task. The larger margins on program-type tasks are more informative precisely because they occur in settings where a local multi-gene representation is more plausible.

Explicit graph and pathway models address a related problem from another direction. P-NET, DCell and MOGONET inject curated biological structure into the model or graph construction [30, 31, 32]. Such methods can avoid the alignment problem of unsupervised layouts: genes that interact biologically may not be close under marginal co-expression [33]. A rigorous next comparison should therefore evaluate 3D voxel carriers, 2D carriers, tuned tabular baselines and explicit graph/pathway models on identical splits and gene sets.

### 4.4 Limitations

Several limitations define the scope of the conclusions. First, the geometric argument does not quantify the size of the stress gap or guarantee that the preserved neighborhood is relevant to a given target. Second, a 3D grid introduces a resolution–sparsity trade-off: increasing dimensionality can reduce flattening distortion but also increases sparsity, computation and sample-complexity demands. Third, the breadth of the program-type evaluations could be extended. GDSC and LINCS already probe drug-response and perturbation mechanism-of-action settings; the drug-response conclusions would benefit from confirmation in larger cohorts with cell-line- or compound-disjoint splits. Fourth, the current sparse 3D backbone is trained from scratch; the comparison to 2D image routes may change if large-scale expression pretraining becomes available for 3D carriers. Fifth, occlusion attribution provides a traceable model-derived ranking, but it does not establish causal gene function.

### 4.5 Future directions

Four directions extend this work naturally. The first is a direct benchmark against explicit biological-graph models on the same tasks. A useful design compares vDeepInsight, matched 2D carriers, tuned tabular baselines, Reactome- or STRING-based graph neural networks, and pathway-sparse networks under identical train/test splits and gene panels, separating the value of unsupervised expression-derived locality from the value of curated biological relations.

A second direction broadens the program-task evaluation. Drug-response cohorts such as CTRP, PRISM and additional GDSC releases, perturbation resources beyond the 20-class LINCS subset, and single-cell perturbation datasets would test whether the GDSC and LINCS patterns generalize across compounds, cell contexts and measurement platforms; compound-family holdout and cell-line-disjoint splits are especially informative here.

A third direction is pretraining for sparse 3D carriers, with candidate objectives including masked-voxel reconstruction, contrastive learning across biological replicates or perturbation pairs, and multi-task prediction across public expression compendia. The informative question is not only whether pretraining improves vDeepInsight, but whether it shifts the balance between 3D carriers and pretrained 2D image backbones.

A fourth direction makes the layout itself more biologically constrained. The current coordinates are derived mainly from expression similarity; hybrid layouts that combine co-expression with pathway membership, protein-interaction networks or regulatory priors offer a route to reducing the alignment problem, paired with ablations that separate the benefit of curated priors from the benefit of the 3D carrier. The synthetic framework used here provides a controlled starting point for those ablations.

## 5 Conclusion

vDeepInsight provides an injective 3D voxel carrier for tabular features, together with a matched framework for evaluating carrier dimensionality; gene expression is the validation setting used here. The four linked evaluations give a consistent result: the 3D layout improves neighborhood fidelity, a sparse local convolution can exploit that fidelity when the target requires it, and under matched evaluation the carrier matches or exceeds the strongest tuned baseline on every task while consistently exceeding the matched 2D carriers. The size of the downstream gain tracks task structure—small on near-saturated marker-type classification, where individual genes and low-dimensional module summaries already carry much of the label, and largest on program-type outcomes. The contribution is a carrier design and evaluation framework showing that a richer spatial carrier preserves neighborhood structure more faithfully and improves prediction when that structure is informative. The carrier places arbitrary tabular features and is not specific to gene expression; all evaluations in this paper use gene-expression data, so the conclusions are stated for that setting, and other structured-feature domains are left to separate work.

## Data availability

The datasets used in this study are publicly available from the Genomic Data Commons/TCGA, the TCGA PanCancer immune-landscape resource, GTEx Portal, DepMap Portal, GDSC database and LINCS/CLUE. Dataset-specific preprocessing, split construction and evaluation units are summarized in Table 1.

**Table 1:** Datasets and evaluation endpoints. Rows are grouped by the role of each analysis. The same split, gene panel, preprocessing and metric were used for all carriers within a matched block. Class counts and split details are given in the text.

| Analysis role | Dataset | Evaluation unit | Primary endpoint |
| --- | --- | --- | --- |
| Carrier fidelity | TCGA, 6,000 genes | 5 carriers, shared genes | Trustworthiness, stress |
| Synthetic capability | Simulated profiles | Signal × layout × kernel | Macro-F1 |
| Marker classification | TCGA / GTEx / CCLE | 5 seeds × 3 panels | Macro-F1 |
| Drug response | DepMap-GDSC, 676 drugs | 3 disjoint splits | Pearson, Spearman |
| Immunogenomic context | TCGA, 17 targets | Raw + residualized | Pearson |
| Perturbation MoA | LINCS L1000, 20 classes | Signature-level split | Macro-F1 |
| Attribution | TCGA models + voxel map | Seed-averaged occlusion | Gene/pathway readout |

**Table 2:** Cross-task performance summary. The table reports the primary endpoint for the 3D voxel carrier and the strongest recorded non-3D baseline in each task. Values are means over the evaluation units defined in Table 1; the table is intended as a compact summary, not as a statistical test table.

| Task | Primary endpoint | 3D voxel | Strongest recorded baseline |
| --- | --- | --- | --- |
| *Marker-type classification* |  |  |  |
| GTEX tissue (51 classes) | Macro-F1 | 0.959 | 0.948 (XGBoost) |
| CCLE lineage (26 classes) | Macro-F1 | 0.754 | 0.709 (XGBoost) |
| TCGA cancer type (33 classes) | Macro-F1 | 0.943 | 0.936 (XGBoost) |
| *Program-type prediction* |  |  |  |
| GDSC drug response (676 drugs) | Per-drug Pearson $r$ | 0.470 | 0.446 (HGB) |
| TCGA immunogenomic-context (17 targets) | Per-target Pearson $r$ | 0.698 | 0.641 (RF) |
| LINCS MoA (20 classes) | Macro-F1 | 0.577 | 0.561 (MLP) |

## Code availability

Code to construct the carriers, train models, run baselines and reproduce the figures will be made publicly available in a GitHub repository after manuscript submission, with an installable package to be released on PyPI.

## Acknowledgements

We thank the Genomic Data Commons/TCGA, GTEx, DepMap, GDSC and LINCS/CLUE projects and their data contributors for making the gene-expression and drug-response datasets used in this study publicly available.

## Funding

This work was partly funded by JSPS KAKENHI Grant Numbers 24K15175, 25KJ1104, JP20H03240 and JP25K02261, Japan and JST CREST Grant Number JPMJCR2231, Japan.

## Competing interests

The authors declare no competing interests.

## Supplementary notes

### Supplementary Note 1. Study design and reproducibility framework

The study was designed as a matched carrier-dimension evaluation. Within each experimental block, the dataset split, selected genes, preprocessing, labels and evaluation metric were fixed before changing the carrier or the baseline model. This design reduces the risk that an apparent 3D advantage is caused by a different split, gene panel or preprocessing path. The evaluation has four components: representation-quality analysis, synthetic neighborhood controls, supervised benchmarks and attribution through the injective voxel map. The project code is organized to reproduce each component and regenerate the figures from the recorded result tables, so figure typography and layout can be revised without re-running model training.

### Supplementary Note 2. Baseline models and parameter selection

Baseline models were selected to test whether the voxel carrier adds information beyond standard tabular and image-carrier alternatives, not to create weak comparators. All tabular baselines used the same training split, selected genes and standardized expression matrix as the corresponding voxel run. Hyperparameter selection was restricted to the training or validation data within each matched block; test labels were not used for model selection.

For marker-type classification, the manuscript baseline set comprised random forests, histogram gradient boosting, XGBoost and the FT-Transformer. The random forest classifier uses 400 trees with class-balanced subsampling when class weighting is enabled. Histogram gradient boosting uses early stopping, validation-fraction monitoring and a leaf-count cap; for larger datasets the implementation applies a memory-aware depth/iteration cap. XGBoost uses histogram-tree construction with a multiclass log-loss objective. The FT-Transformer baseline uses a reduced tabular transformer configuration with 256 input features, token dimension 64, eight attention heads, two layers, dropout 0.1, Adam optimization and early stopping through a fixed epoch budget. These settings were kept as recorded defaults rather than re-tuned post hoc for individual figures.

For GDSC drug-response regression, the tabular regressors were fitted per drug on the cell lines with observed response for that drug. The random forest regressor used 200 trees; histogram gradient boosting used 200 boosting iterations, depth 6, learning rate 0.1 and early stopping; ridge regression used *α* = 1.0. The reported endpoint is the held-out per-drug correlation between predicted and observed ln IC50, summarized over the 676 drugs and three cell-line-disjoint splits.

For neural carriers, the mainline 3D model used the plain sparse-convolutional backbone with multi-scale mean–max pooling. Classification runs used focal loss with label smoothing and a capped class-weight term, as specified in the recorded run flags; regression runs used a smooth-L1 objective. The 2D image carriers used compact 2D CNN consumers trained from scratch under the same split, gene-panel and evaluation protocol as the 3D carrier. No ImageNet-pretrained, task-pretrained or large-scale vision backbone was used. The 2D models therefore serve as matched carrier controls for isolating the effect of carrier dimensionality and assignment geometry, rather than as reproductions of optimized DeepInsight-2D pipelines reported in previous studies. A deeper residual sparse-3D backbone was implemented as an ablation but did not provide consistent improvement, so it was not used as the headline configuration.

## Supplementary figures

**Figure S1:**
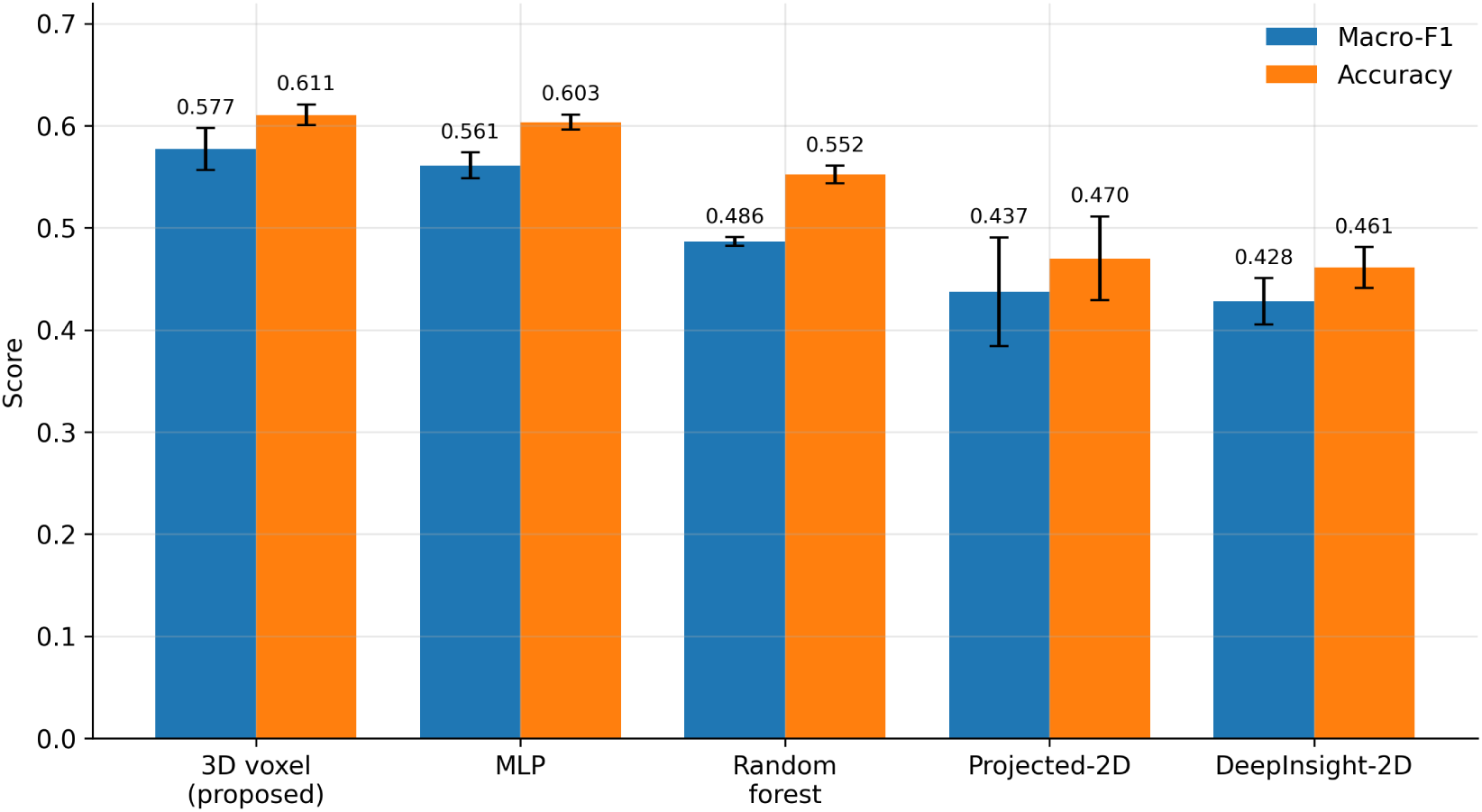
LINCS mechanism-of-action classification. Supplementary Figure S1 reports performance on the 20-class LINCS L1000 MoA task under a signature-level split. The two panels show macro-F1 and accuracy by method, with bars denoting mean performance over seeds and error bars denoting SD. The 3D voxel carrier uses the plain multi-scale mean–max sparse-CNN backbone used throughout the main analyses and attains the highest macro-F1 and accuracy among the recorded methods.

**Figure S2:**
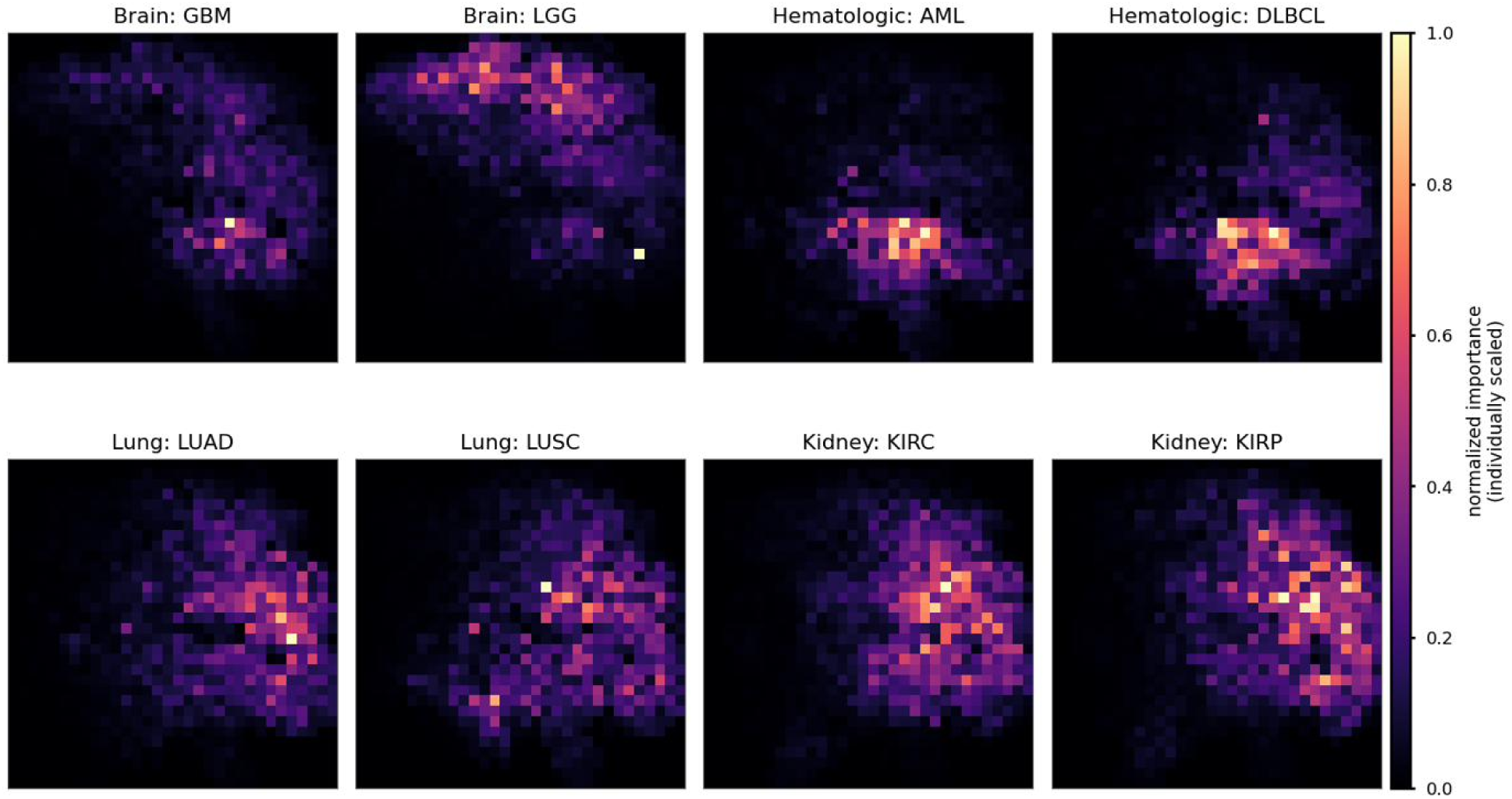
Per-class attribution overview (TCGA). Supplementary Figure S2 shows two-dimensional projections of seed-averaged 3D voxel occlusion attribution for eight representative TCGA cancer types grouped into four related tumor groups: brain (GBM, LGG), hematologic (AML, DLBCL), lung (LUAD, LUSC) and kidney (KIRC, KIRP). Importance is min–max scaled within each cancer-type panel so that the spatial pattern of each class can be compared visually. In every class, attribution is concentrated in a limited number of compact layout regions rather than spread uniformly across the carrier, and the active regions differ across cancer types and related subtype pairs. Because the voxel-to-feature assignment is one-to-one, each highlighted region maps to a defined set of genes; these maps provide the spatial, gene-resolved basis for the class-specific attribution summaries in Figures S4 and S5.

**Figure S3:**
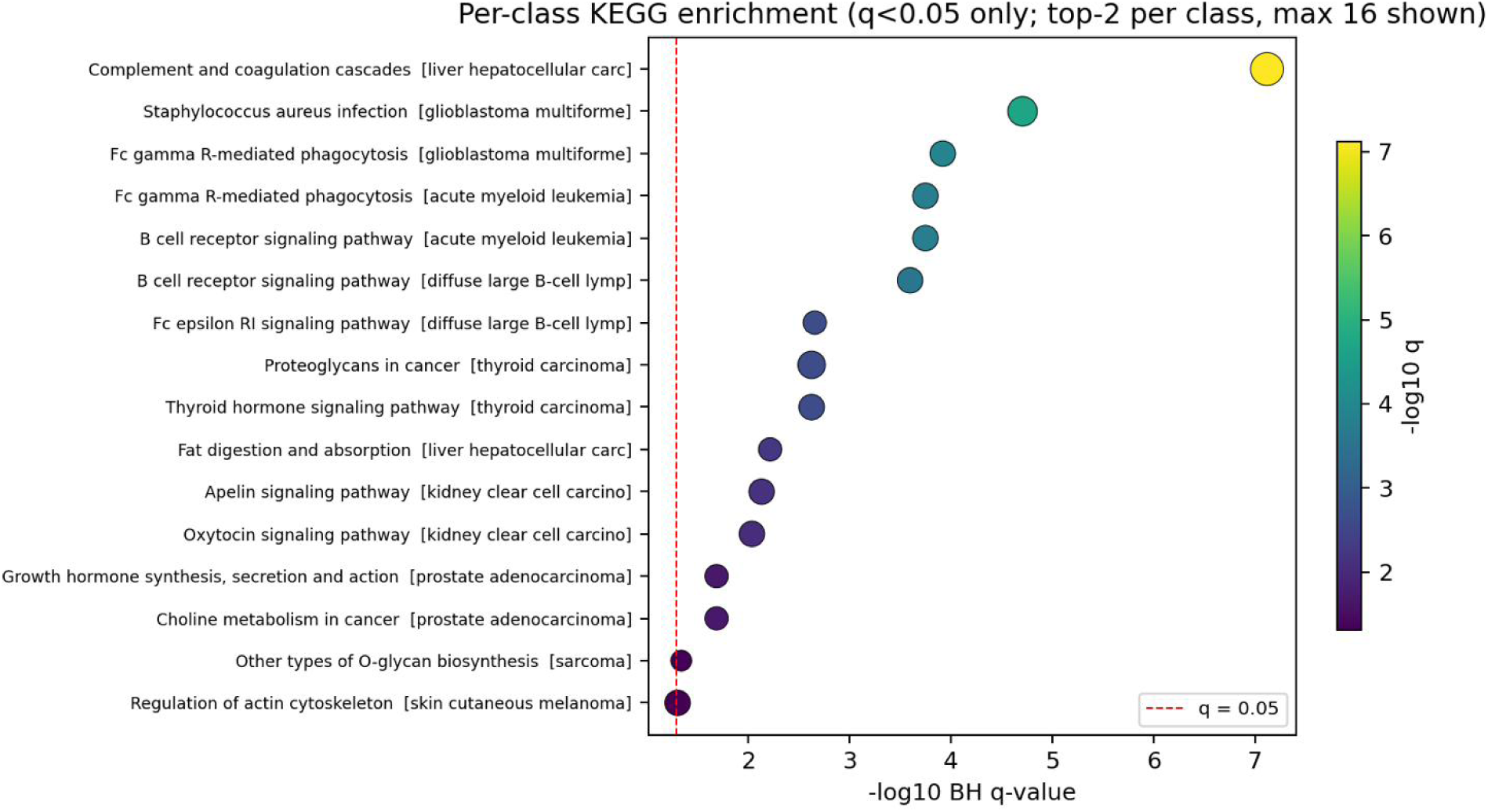
Per-class pathway enrichment (TCGA). KEGG pre-ranked enrichment of seed-averaged gene-level attributions by cancer type. The maps summarize the functional programs associated with the genes used by the trained model for each class. Leading enriched pathways are consistent with known biology of the corresponding tumor types and provide pathway-level functional interpretation of the gene-resolved attribution results.

**Figure S4:**
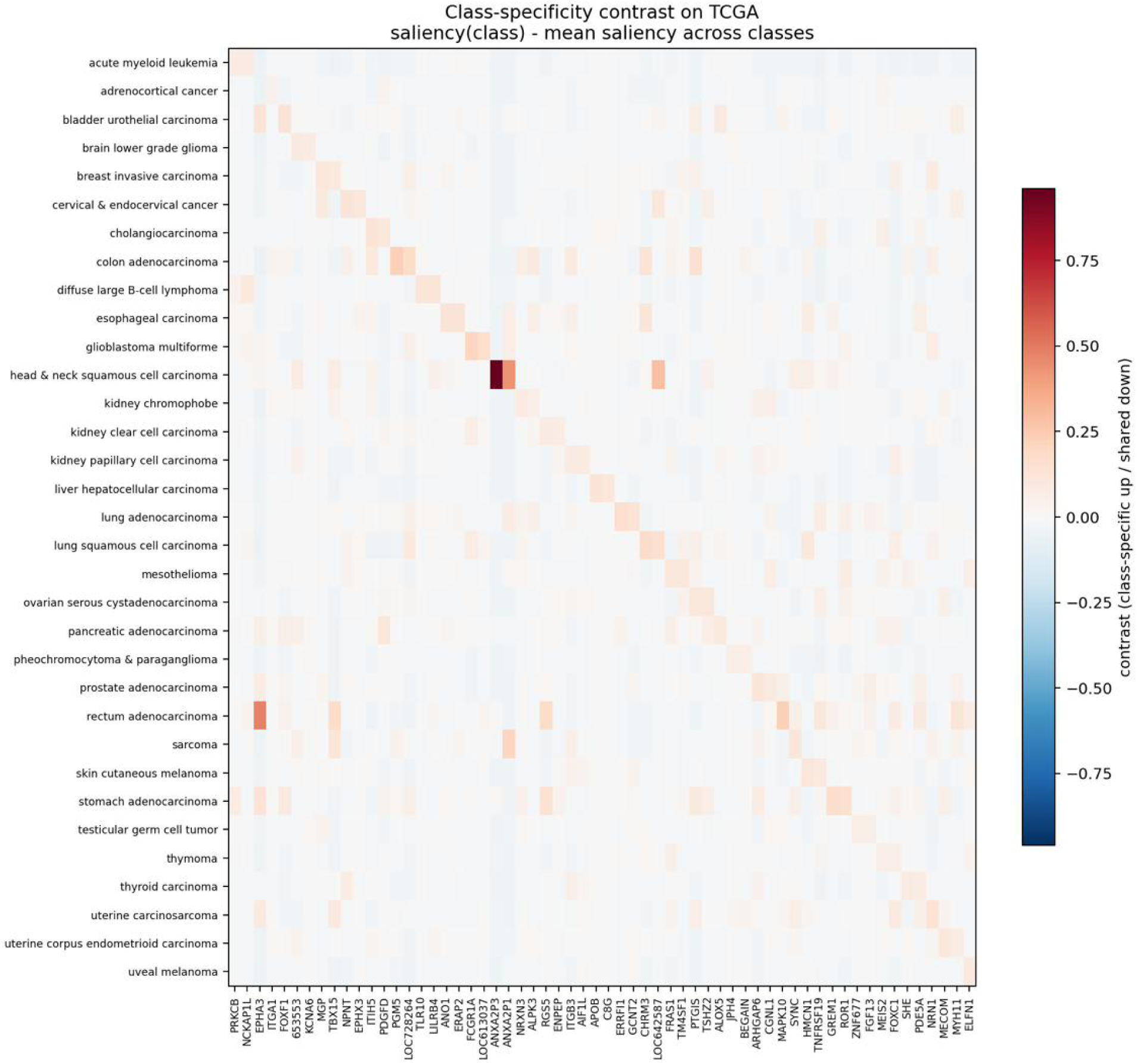
Class specificity of attributions (TCGA). Comparison of attribution profiles across cancer types, showing that leading genes are largely class-specific rather than shared across all classes.

**Figure S5:**
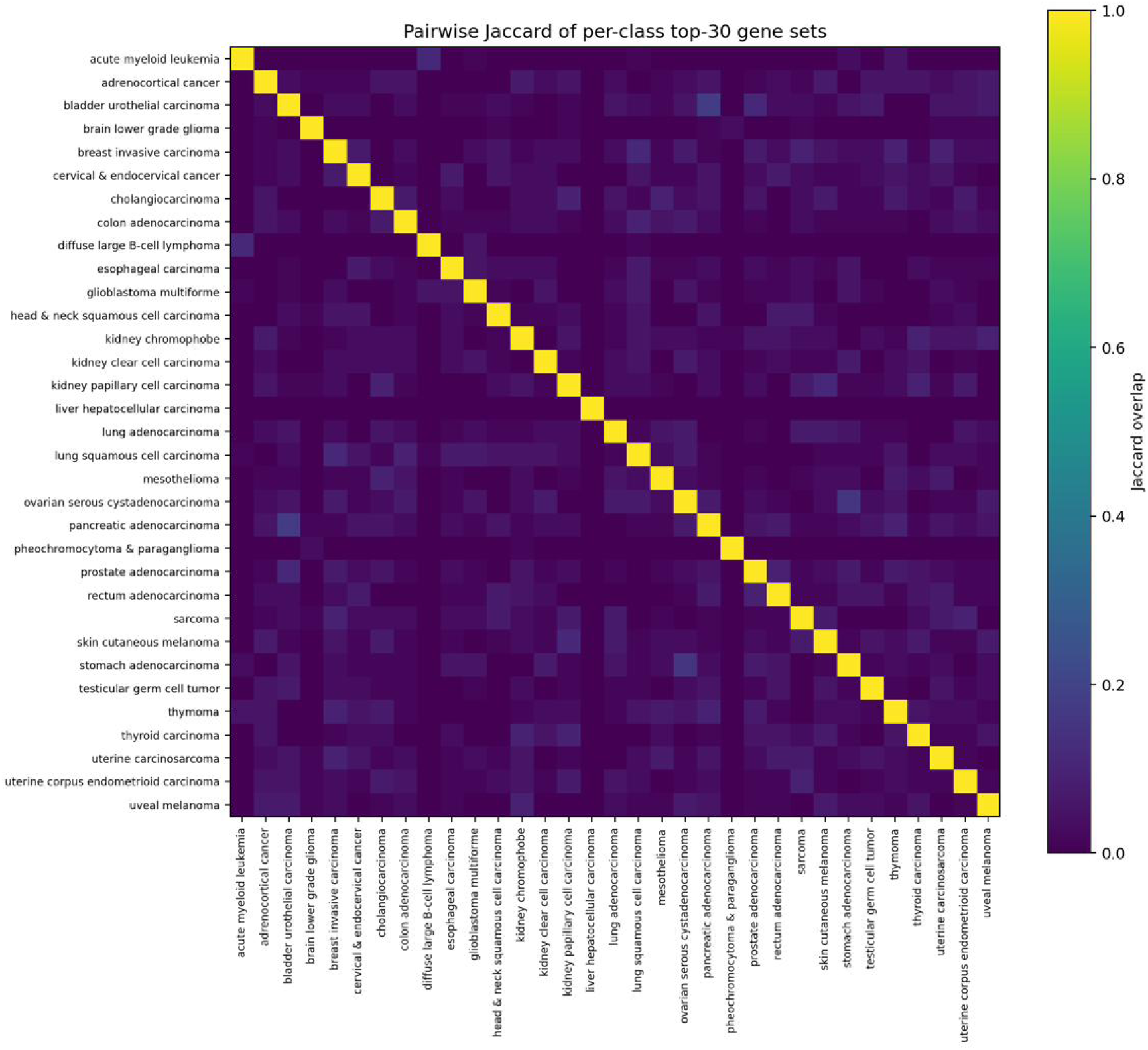
Cross-class distinctness of top-gene sets (TCGA). Pairwise Jaccard overlap between per-class top-attributed gene sets. Low off-diagonal overlap indicates that different cancer types rely on largely distinct gene sets, complementing the spatial attribution overview and pathway-level functional summaries.

## Notes

### Competing Interest Statement

The authors have declared no competing interest.

